# A novel fully-spliced accessory gene in equine lentivirus with distinct Rev-responsive element

**DOI:** 10.1101/2022.01.30.478423

**Authors:** Xiangmin Zhang, Jiwei Li, Mengmeng Zhang, Bowen Bai, Weiwei Ma, Yuezhi Lin, Xing Guo, Xue-Feng Wang, Xiaojun Wang

**Author notes:** Corresponding author. (X-FW); (XJ). **Author Contributions Conceptualization:** Xiangmin Zhang, Xue-Feng Wang, Xiaojun Wang. **Funding Acquisition**: Xue-Feng Wang, Xiaojun Wang. **Investigation**: Xiangmin Zhang, Jiwei Li, Mengmeng Zhang, Bowen Bai, Weiwei Ma, Xing Guo, Xue-Feng Wang **Methodology**: Xiangmin Zhang, Jiwei Li. **Formal analysis**: Xiangmin Zhang, Xue-Feng Wang, Yuezhi Lin. **Supervision**: Xue-Feng Wang, Xiaojun Wang. **Validation**: Xiangmin Zhang, Jiwei Li, Mengmeng Zhang, Weiwei Ma. **Writing-Original Draft Preparation**: Xiangmin Zhang, Xue-Feng Wang, Xiaojun Wang. **Writing-Review & Editing**: Xiangmin Zhang, Xue-Feng Wang, Xiaojun Wang.

## Abstract

All lentiviruses encode an accessory protein Rev, whose main biological function is to mediate the nuclear export of unspliced and incompletely spliced viral transcripts by binding to a viral cis-acting element (termed the Rev-responsive element, RRE) that is located within env-encoding region. Equine infectious anemia virus (EIAV) is a member of the Lentivirus genus in the Retroviridae family, and is considered an important model for the study of lentivirus pathogenesis. Here, we identified a novel transcript from the EIAV genome that encodes a viral protein, named Mat, with unknown function. The transcript *mat* is fully spliced and comprises parts of the coding regions of MA and TM. Interestingly, the expression of Mat depends on Rev and the CRM1 pathway. Rev could specifically bind to Mat mRNA to promote its nuclear export. We further identified that the first exon of Mat mRNA, which is located within the Gag-encoding region, acts as an unreported RRE. Altogether, we identified a novel fully spliced transcript *mat* with an unusual RRE, which interacts with Rev for nuclear export through the CRM1 pathway. Our findings may help to expend the understanding of gene transcription and expression of lentivirus.

**Author summary:** In lentiviruses, the nuclear export of viral transcripts is an important step in controlling viral gene expression. Generally, the unspliced and incompletely spliced (that is, intron-containing) transcripts are exported via the CRM1-dependent export pathway in a process mediated by the viral Rev protein by binding to the Rev-responsive element (RRE) located within the Env-coding region. However, the completely spliced (intronless) transcripts are exported via an endogenous cellular pathway which is Rev independent. Here we identified a novel intronless transcript from EIAV and demonstrated that it encodes a viral protein, termed Mat. Interestingly, we have determined that the expression of Mat depends on Rev, and identified that the first exon of Mat mRNA could specifically bind to Rev and be exported to the cytoplasm, which suggests that first exon of Mat mRNA is a second RRE of EIAV. These findings update the EIAV genome structure and highlight the diversification of post-transcriptional regulation patterns in EIAV, and provides important insights into the Rev-dependent nuclear export of completely spliced transcripts in lentiviruses.

## Introduction

Equine infectious anemia virus (EIAV) is a member of the Lentivirus genus of the Retroviridae family and primarily infects *equids.* Unlike other lentiviral infections characterized by chronic degenerative progression, EIAV infection in horses usually leads to an acute disease course [1]. Moreover, after approximately one year of infection, most infected animals are able to control viral replication and progress to inapparent carriers [2]. An inapparent carrier can still recrudesce disease under stress or immune suppression [1, 3]. These phenomena indicate that infected horses have gained immunologic control over the EIAV. Thus, the EIAV system provides a good model for the study of lentivirus replication and lentiviral vaccine development [4].

Lentiviruses, including the human immunodeficiency virus type 1 (HIV-1), simian immunodeficiency virus (SIV) and EIAV, like all retroviruses, are RNA viruses that replicate via a DNA intermediate (called proviral DNA) which is integrated into the host genomic DNA. In addition to the three structural proteins (Gag, Pol and Env) common to retroviruses, the lentiviral genome also encodes several accessory proteins. Current knowledge indicates that EIAV encode at least four accessory proteins (Tat, S2, Rev and Ttm) [5], while primate lentiviruses (HIV-1 and SIV) encode six accessory proteins (Tat, Rev, Nef, Vpu, Vif, Vpr/Vpx) [6, 7]. These accessory proteins are not only directly involved in viral transcriptional and posttranscriptional regulation (Tat and Rev) [8], but also regulate various stages of viral replication by interacting with host proteins (e.g., Nef, Vpu, Vif, Vpr, S2) [9–11]. At both ends of the lentiviral genome are identical long terminal repeats (LTR) [12]. As a viral promoter, 5’LTR regulates viral genome transcription by binding with host RNA polymerase II, other host proteins (such as cyclinT) as well as virus-encoded proteins including Tat [8, 13].

The lentiviral proviral DNA acts as a transcription template to synthesize viral mRNA in the nucleus. Transcription is initiated at the U3-R border of the 5’LTR and terminates at the R-U5 border of the 3’LTR [14]. A full-length genome mRNA is first transcribed, which is a primary unspliced transcript, and then most of this transcript undergoes alternative splicing to generate multiple spliced transcripts, allowing expression of several viral proteins. In HIV-1, more than 40 transcripts have been found, which are responsible for the expression of the viral proteins [8]. In EIAV, five transcripts have been identified in earlier studies [5, 15, 16], including an unspliced full-length mRNA transcript encoding Gag and Pol protein as well as genomic RNA for encapsidation into progeny virions, an incompletely spliced transcript encoding Env and S2 protein, three fully spliced subgenomic transcripts encoding the accessory proteins (Tat, Rev) and a predicted Ttm protein whose function is unknown. Recently, our lab identified an unreported transcript in EIAV encoding the S4 protein, which was found to facilitate viral release by antagonizing the antiviral activity of the host protein tetherin (manuscripts, under review).

In lentiviruses, the synthesis and production of all viral mRNA takes place in the nucleus and the mRNA is then transported to the cytoplasm by different nuclear export pathways [17]. For example, in HIV-1, the unspliced full-length and incompletely spliced transcripts encoding viral structural proteins (Gag, Pol and Env) and some accessory proteins (vif, vpr and vpu), are exported to the cytoplasm utilizing the CRM1-dependent export pathway mediated by the viral Rev protein. However, all the completely spliced transcripts encoding the viral accessory proteins Tat, Rev and Nef, are exported to the cytoplasm by an endogenous cellular pathway used by cellular mRNAs independent of Rev [8, 18]. Other lentiviruses, such as EIAV and FIV, also utilize the Rev-mediated CRM1 pathway to export incompletely spliced mRNA transcripts [19]. The Rev protein is an accessory protein of all lentiviruses, and interacts with a specific cis-acting element termed the Rev responsive element (RRE) at the viral pre-mRNA [20, 21]. The lentiviral RRE is a highly structured viral RNA sequence that is generally located in the Env-coding region [20]. Although the lentiviral RREs lack sequence homology, they may share specific stem-loop structures. Some studies have identified that the RRE in HIV-1 is about 350 nt and that in EIAV is about 555 nt [20, 22]. The molecular details of the Rev-mediated incompletely-spliced mRNA export have been well characterized [23]. A single Rev molecule first binds to RRE, and subsequently multiple Rev molecules undergo a series of Rev-Rev oligomerizations to form a multimer, which subsequently recruits the nuclear export factor CRM1 to promote the export of the incompletely spliced mRNA into the cytoplasm. Thus, Rev binding and subsequent multimerization are necessary for Rev-RRE complex function. Previous studies have identified the RNA binding domain and multimerization of EIAV Rev [24–27]. Considering that Rev-mediated RNA export is essential for lentiviral replication, the Rev-RRE complex is a potential therapeutic target in lentiviruses.

In the present study, we discovered a novel viral protein in EIAV and named it Mat. The protein is translated from an unreported fully-spliced transcript from the EIAV genome. Interestingly, the expression of Mat depends on the Rev-mediated RNA nuclear export pathway. We provide evidence that Rev binds to Mat mRNA and is required for the nuclear export of Mat mRNA.

## Results

### Discovery and identification of a novel protein-encoding gene from EIAV

According to the genome structure and transcriptional characteristics of EIAV, we designed an upstream primer (p399), and two downstream primers (p8144 and p8081), as shown in Fig 1A. The PCR products were cloned and sequenced, and the sequences were then aligned to the EIAV genome (GenBank accession numbers: GU385361.1). In addition to the S4 and S5 transcripts that have been described in other manuscripts (under review), a novel EIAV mRNA with single-splicing event was also discovered by sequence alignment. The mRNA utilized a 5’ splice donor site (SD) at nt863 (^862^TG^863^|^864^GG^865^) and a 3’ splice acceptor site (SA) at nt7850 (^7848^AT^7849^|^7850^AT^7851^). The transcript includes a 447 bp open reading frame (ORF), and contains two exons, with the first exon (393 bp) partially overlapping with the MA subunit coding region of the Gag protein (in the same reading frame) and the second exon (54 bp) partially overlapping the 3’terminal region of the TM coding region of the Env protein (in a different reading frame). We named the novel encoding gene *mat* and its encoding protein Mat. According to the splicing pattern of the mat transcripts, we designed specific nested PCR primers (forward primers, p555 and p643; reverse primers, p7878 and p7855) to amplify part of this transcript sequence (about 250bp) from multiple tissues (lymph gland, testis, liver, kidney, brain, spleen, marrow and heart) of horses infected with EIAV_LN40_ (Fig 1B). The obtained PCR products were sequenced and the junction between the SD^863^ and SA^7850^ sites, namely mat-specific splicing sites was found. We also amplified the mat-specific bands from the equine monocyte-derived macrophages (eMDM) infected with EIAV_DLV121_ and EIAV_DLV36_ *in vitro* (Fig 1C). All these data suggested that there were mat transcripts in tissues or cells infected with EIAV. Furthermore, to detect the expression of Mat in EIAV-infected cells, we generated a mouse monoclonal antibody against the peptide (ENAKSSYISCNNASI) corresponding to the translated product of the second exon of mat. Using this antibody, we detected an about 20-kDa specific band in eMDM cells infected with EIAV_DLV121_ and EIAV_DLV36_ but not in uninfected cells (Fig 1D), which suggests that the Mat protein is expressed in EIAV-infected cells.

**Fig 1.**
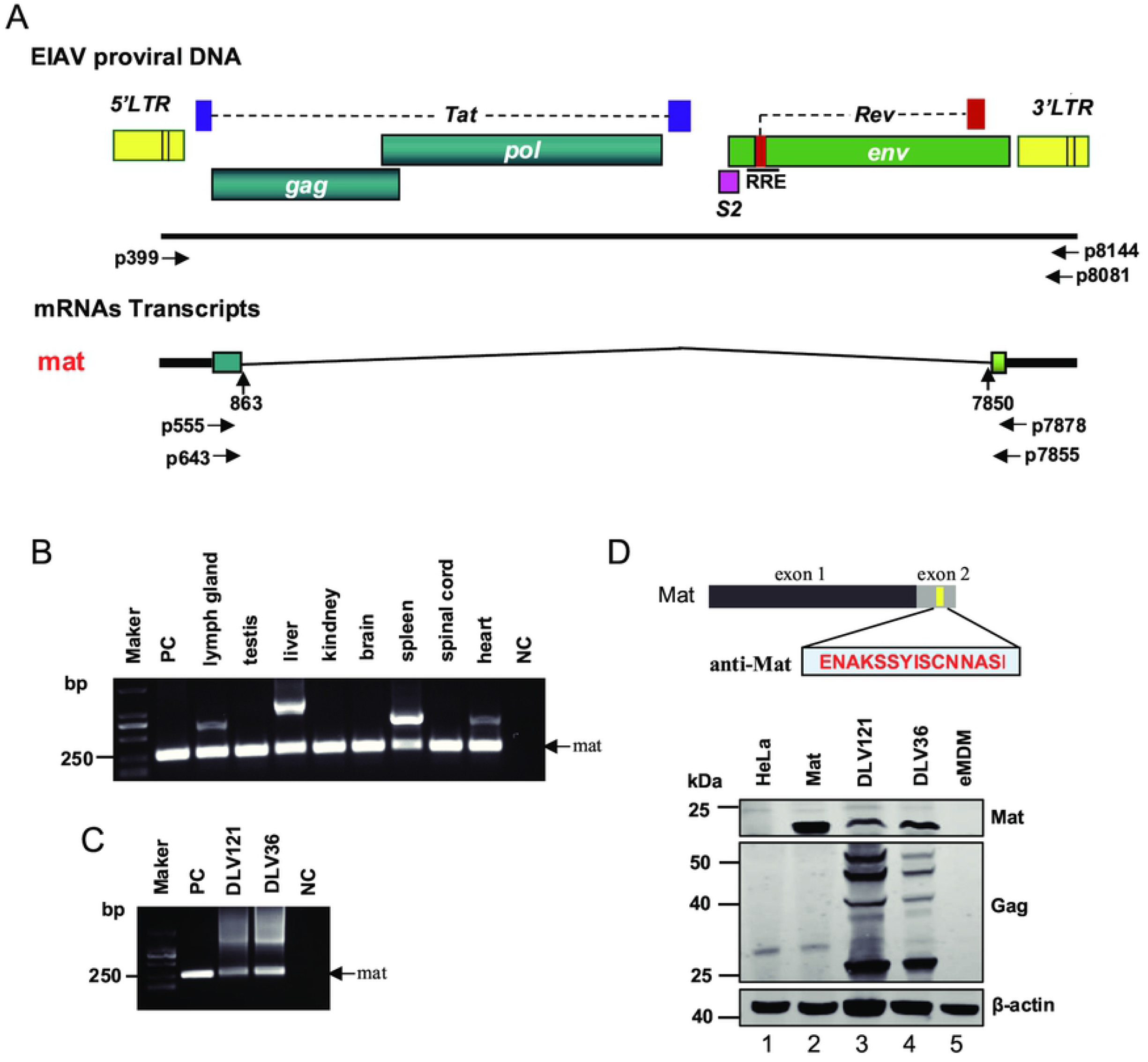
Discovery and identification of a novel protein-encoding gene from EIAV. (A) A schematic representation of the EIAV genome is displayed at the top of the figure. Known ORFs are shown as boxes with different colors. The position of RRE is depicted with a heavy line. The splicing pattern of mat transcripts is shown below the genomic structure. Rectangles represent exons of genes. Vertical arrows represent splice site of mRNA. Horizontal arrows show the locations of oligonucleotide primers for amplification of cDNAs. (B) Identification of the mat gene from multiple tissues (lymph gland, testis, liver, kidney, brain, spleen, marrow and heart) of horses infected with EIAV_LN40_. PC, mat sequence inserted into pMD18T-vector as a positive control. NC, negative control. (C) Identification of the mat gene from eMDM cells infected with EIAV_DLV36_ and EIAV_DLV121_. (D) Identification of Mat from eMDM cells infected with EIAV_DLV36_ and EIAV_DLV121_. The Mat-specific antiserum (described in Materials and Methods) was used for the immunodetection of Mat protein. An anti-EIAV p26 antiserum was used to monitor the effect of infection. Hela cells transfected with empty plasmids were used as a negative control (lane 1). Hela cells transfected with a codon-optimized Mat expression plasmid were used as a positive control (lane 2). eMDM cells isolated from healthy horses were used as a specific control (lane 5).

### The expression of Mat depends on the nuclear export activity of Rev

To express the novel protein Mat in a eukaryotic expression system, its coding sequences were inserted into a pcDNA3.1 vector. We were unable to detect the expression of Mat when transfected HEK239T cells with the Mat expression plasmid alone. However, when HEK293T were cotransfected Mat with the Rev expression plasmid, Mat proteins could be observed clearly using western blotting (WB) (Fig 2A). This Mat expression pattern was also observed in fetal donkey dermal (FDD) cells (Fig 2B), which are permissive cells for the replication of EIAV [28, 29]. GFP was used here as a specific control to monitor transfection efficiencies. This result suggested that Rev is necessary for the expression of Mat. Furthermore, when HEK293T cells were transfected with different doses of Rev, the expression of Mat increased in a dose-dependent manner (Fig 2C).

**Fig 2.**
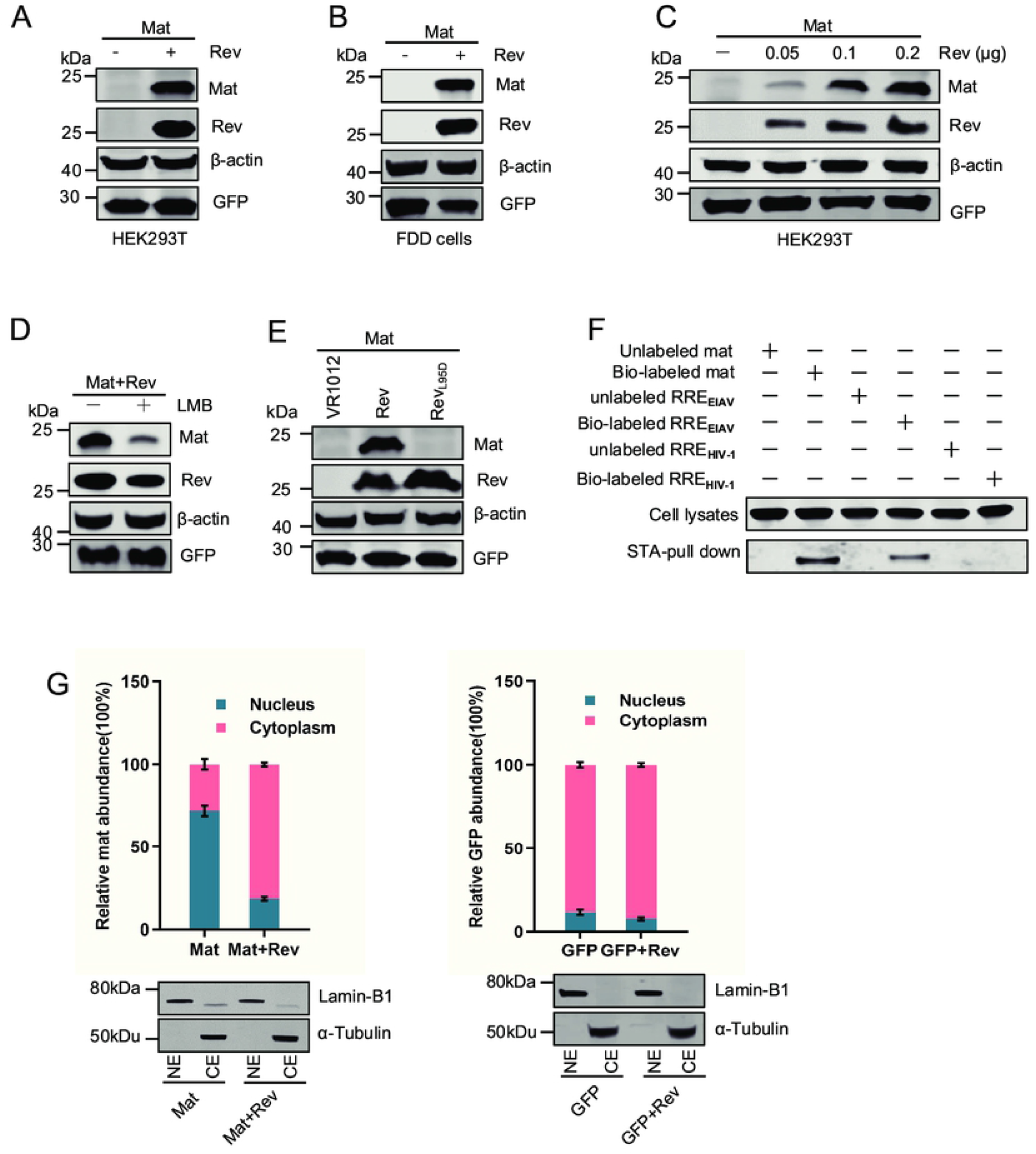
The expression of Mat depends on Rev. A Mat constructs was transiently cotransfected into HEK293T cells (A) or FDD cells (B) with Rev expression plasmids or empty plasmids. Mat expression was analyzed with WB using a mouse anti-HA antibody 48h post transfection (hpt); β-actin expression was detected using an actin-specific antiserum to monitor unity of blotting and equal loading of the gel; GFP expression was quantified using a GFP-specific antiserum to monitor transfection efficiencies. (C) An Mat constructs was transfected into HEK293T cells, along with increasing amounts of a Rev expression plasmid. (D) LMB inhibited Rev-independent expression of Mat. HEK293T cells were cotransfected with Mat constructs and Rev expression plasmids. At 12 hpt, cells were treated with leptomycin B (0nM [-] or 50nM [+], as indicated). At 24 h post drug treatment, cells were harvested and subjected to WB. (E) No Mat protein expression was detected when cotransfected with Mat constructs and Rev_L95D_ expression plasmids. Rev_L95D_ is a mutant of Rev without the ability to mediating mRNA export to the cytoplasm. VR1012 was used as an empty vector control. (F) Interaction between mat mRNA and Rev protein was determined using an RNA-pull down assay. Cell lysates from HEK293T transfected with Rev expression plasmids were incubated with biotin-labeled or unlabeled RNA transcribed *in vitro.* The biotin-labeled mat mRNA, EIAV RRE or HIV-1 RRE were pulled down with streptavidin (STA)-conjugated magnetic beads, followed by WB of co-purified proteins using anti-Flag antibody. Levels of pulled-down Rev are shown in a bar chart. (G) Distribution of cytoplasmic and nuclear mat or GFP RNAs in HEK293T cells at 36 hpt was analyzed using qPCR with primers specific to mat or GFP mRNA. Ratios of cytoplasmic to nuclear mRNAs were then calculated and are presented. LaminB1 and α-tubulin were detected using WB to assess the efficacy of cell fractionation. All of the experiments were performed three times, and a representative result is shown.

It is well known that Rev is a nuclear-cytoplasmic shuttling protein that can facilitate the export of viral mRNAs from the nucleus via the CRM1-mediated pathway [8, 21]. To investigate whether the expression of Mat was related to the nuclear export activity of Rev, HEK293T cells were cotransfected with Mat and Rev plasmids and then were treated with Leptomycin B (LMB), which is an inhibitor of CRM1. As shown in Fig 2D, the levels of Mat expression significantly decreased in the presence of LMB. In addition, a Rev mutant (RevL95D), that disrupted nuclear export activity [26, 27], was unable to mediate the expression of Mat to the same extent as the wild-type Rev (Fig 2E). These results indicated that Rev-mediated Mat expression depends on the nuclear export activity of Rev through the CRM1-pathway.

To assess whether mat mRNAs interact with Rev protein, an RNA-pull down assay was performed using biotin-labeled mat RNA and streptavidin (STA) beads. WB analysis with anti-Flag antibody was used to detect whether any Rev protein was present in the cell lysates in the mat mRNA pulldown. The results suggest that EIAV Rev is able to specifically bind to mat and EIAV RRE mRNAs (reported RNA export element in EIAV, used as a positive-binding control) but not to HIV-1 RRE mRNAs (RNA export element in HIV-1, used as a specific-binding control) (Fig 2F). Furthermore, we isolated nuclear and cytoplasmic RNAs from HEK293Tcells after cotransfection with Mat and Rev or empty plasmids, and the RNAs were subsequently analyzed using quantitative PCR (qPCR). Our results showed that most of the mat mRNAs were retained in the nucleus in the absence of Rev, whereas the proportion of mat mRNAs in the cytoplasm increased markedly when cotransfected with Rev (Fig 2G, left). By contrast, the presence of Rev made no difference to the nucleocytoplasmic ratio of GFP mRNAs (Fig 2G, right). These results imply that Rev is able to specifically promote the export of mat mRNAs from the nucleus to the cytoplasm. To evaluate the efficiency of separation, LaminB1 and α-Tubulin protein were used as controls for nuclear and cytoplasmic proteins respectively (Fig 2G, bottom). All these results together demonstrate that the expression of Mat depends on Rev, and Rev is able to promote the transport of mat mRNAs by binding to them.

### The first exon of mat is necessary for Rev-mediated expression of Mat

The ORF of mat contains two exons, with the second one having only 54 nucleotides. In order to map the determinants of Mat expression more precisely, we constructed a Mat-exon1 expression plasmid (Fig 3A). As expected, we found that expression of Mat-exon1 also depends on Rev, which is consistent with the expression characteristics of Mat (Fig 3B). To further confirm this phenomenon in multiple EIAV strains, we constructed 6 mat-exon1 expression plasmids with DNA sequences, each corresponding to the N-terminus of the gag gene coding sequence from one of the different strains, as described in the Materials and Methods. Then we cotransfected these mat-exon1 expression plasmids either with Rev or with an empty plasmid. WB analysis demonstrated that in all cases, Mat-exon1 was expressed only in the presence of Rev (Fig 3C). It further confirmed that Rev is necessary for Mat-exon1 expression in the different EIAV strains. We then used an RNA pull-down assay to assess the interaction between mat-exon1 mRNA and Rev. The results demonstrated that mat-exon1 mRNA can also specifically bind to Rev (Fig 3D). In addition, we separated the cytoplasm and nucleus of the transfected cells and quantified the RNAs in the different fractions separately. The qPCR results showed that accumulation of mat-exon1 mRNAs in the cytoplasm increased in the present of Rev (Fig 3E). These results identify the first exon of mat as being the determinant of Rev-mediated Mat expression, with the first exon demonstrating the same characteristics as the full-length mat mRNAs in terms of protein expression, Rev binding, and nucleocytoplasmic transport of mRNAs.

**Fig 3.**
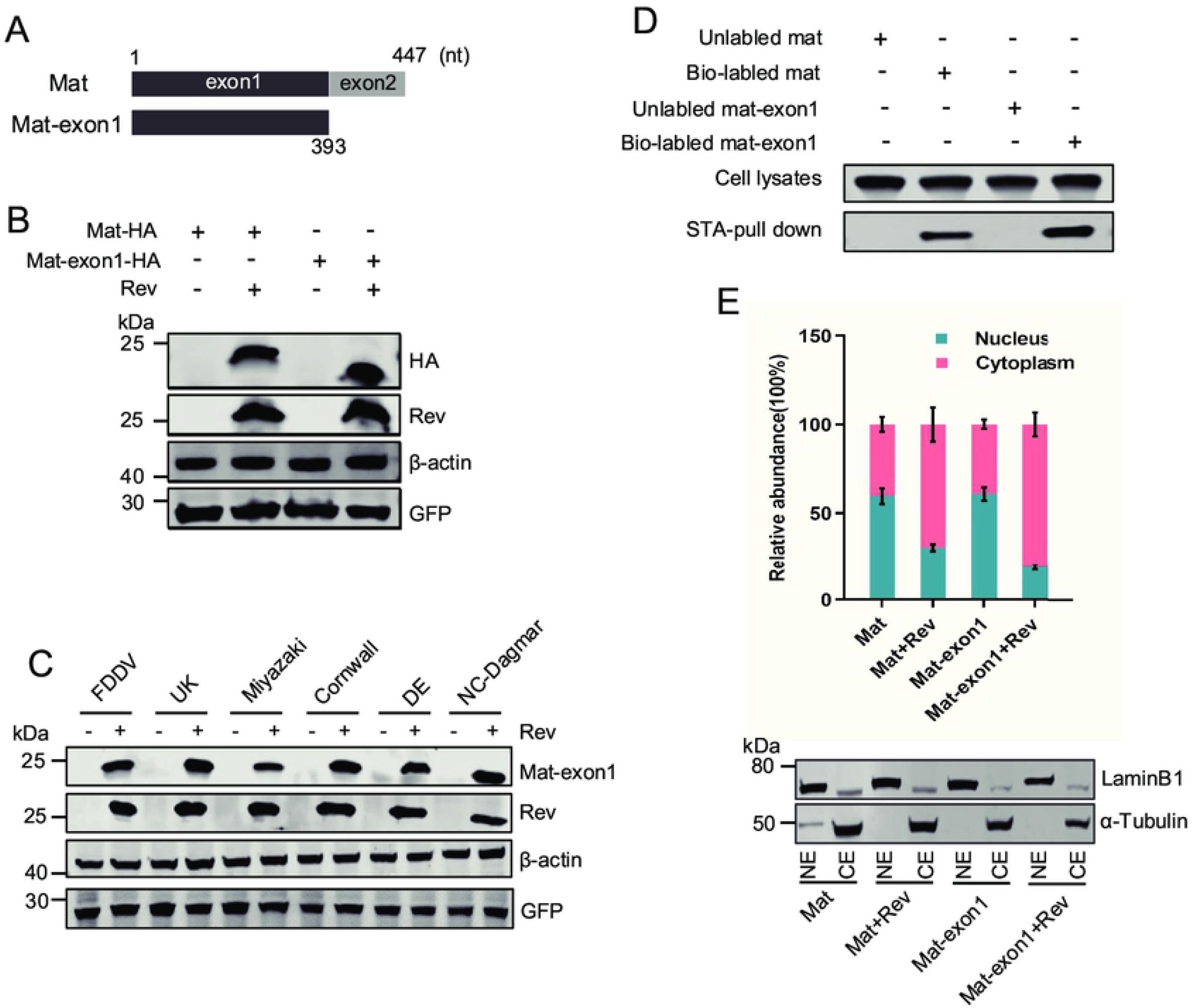
Mapping of the determinants of Rev-mediated expression of Mat. (A) Schematic diagram of the mat ORF coding sequence with 447 nucleotides, which contains two exons (exon-1:1-393nt; exon-2:394-447nt). The exon-1 sequence was inserted into pcDNA-HA to express the Mat-exon1 protein. (B) HEK293T cells were cotransfected Mat or mat-exon1 expression constructs with Rev or empty plasmids. Mat expression was analyzed with WB using a mouse anti-HA antibody; GFP, derived from pEGFP-N1, was quantified using a GFP-specific antiserum to monitor transfection efficiencies. (C) HEK293T cells were cotransfected each of six different Mat-exon1 plasmids, which were derived from six EIAV strains as indicated, with Rev or empty plasmids. Mat-exon1 proteins were detected 48 hpt with WB. (D) An RNA-pull down assay was performed to verify the interaction of mat-exon1 mRNA with Rev protein. The Rev pulled down by mat mRNA acted as a positive control. Biotin-unlabeled mRNAs were used to indicate the specificity of this assay. (E) Distribution of cytoplasmic and nuclear mat-exon1 mRNAs in HEK293T cells at 36 pht was analyzed using qPCR. The subcellar distribution of mat mRNAs was analyzed as a control.

### Different key domains of Rev underlie the mechanism by which the expression of Gag and Mat is regulated

Previous studies have described the four functional domains within Rev: a nuclear export signal (NES), an RNA binding domain (RBD), a nuclear localization signal (NLS) and a functionally unknown non-essential domain (ND) [30]. In order to map the key domain responsible for the Rev mediation of Mat expression, we constructed five Rev mutants, including the aforementioned four mutants with defects in the respective functional domains [31], and a multimerization-defective mutant Rev_L95D_ that lost the capacity to promote export of the gag-pol mRNAs from the nucleus to the cytoplasm [27] (Fig 4A). A Mat expression vector was cotransfected into HEK293T cells with each Rev mutant separately. After 48 h, the level of Mat protein was assessed using WB and the distribution of mat mRNAs in the cytoplasm and nucleus was analyzed using qPCR as the ratio of nuclear to cytoplasmic mat mRNA. The WB analysis results showed that only the Rev_WT_ and Rev_mNES_ were able to mediate the expression of Mat, while the other four mutants almost completely lost this function (Fig 4B). The qPCR analysis also showed that mat mRNAs were mainly distributed in the cytoplasm in the presence of Rev_WT_ or Rev_mNES_, while the other Rev mutants showed results similar to the absence of Rev, where mat mRNA was mainly distributed in the nucleus (Fig 4C). These results indicate that the RBD, NLS and ND domains of Rev and their multimerization play an important role in mediating the nuclear export of mat mRNAs.

**Fig 4.**
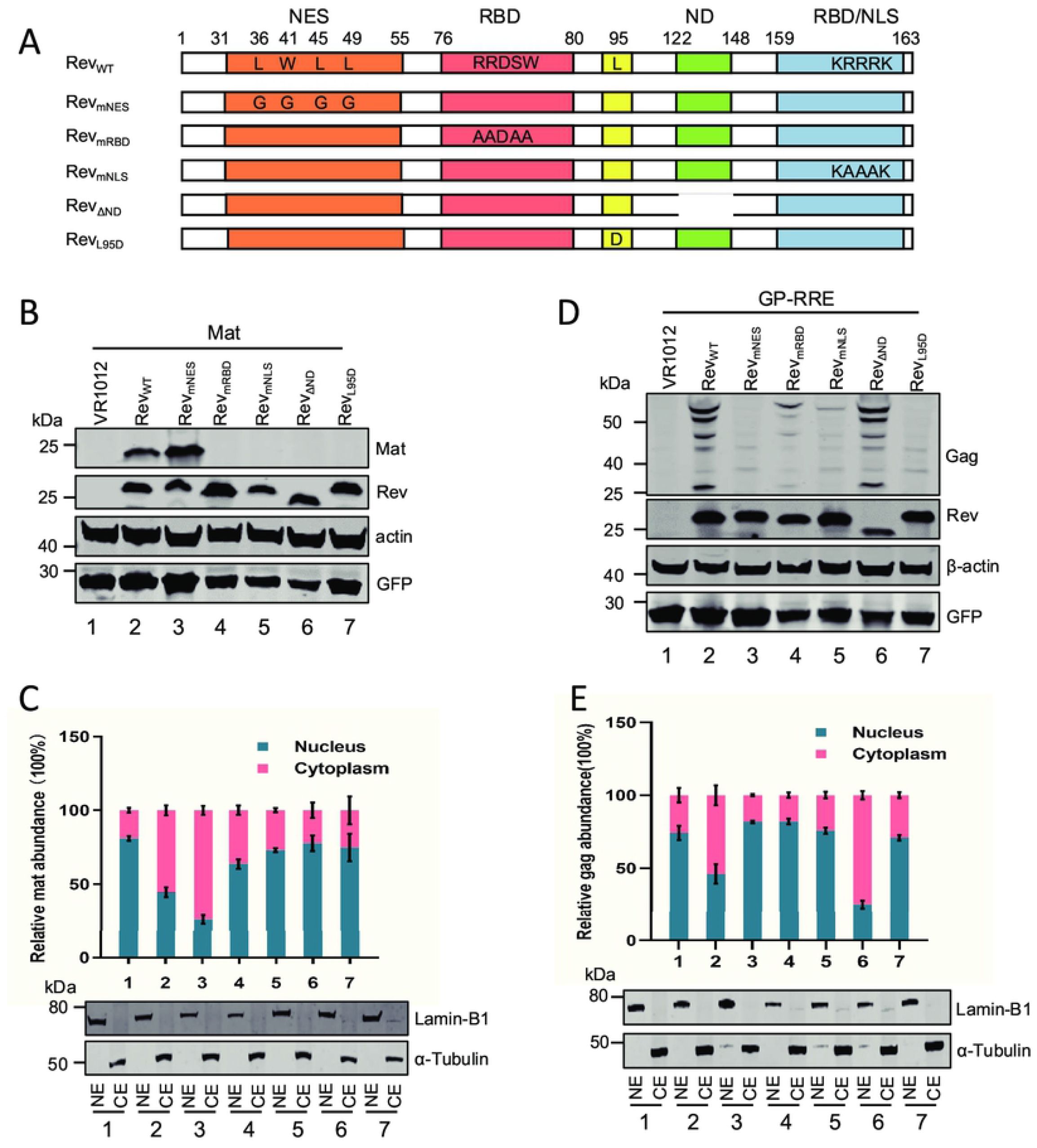
The key domains of Rev mediating the expression of Gag and Mat are different. (A) Schematic diagram of wild-type EIAV Rev, which contains four functional domains: NES (nuclear export domain; aa 31-55); RBD (RNA-binding domain; aa76-88 or aa159-163); ND (Non-essential domain; aa122-148); NLS (nuclear localization domain; aa159-163). The structures of Rev five mutants are: Rev_mNES_, the mutation of Rev from L36, W41, L45, L49 to G; Rev_mRBD_, the mutation of Rev from RRDSW to AADAA; Rev_mNLS_, the mutation of Rev from KRRRK to KAAAK; Rev_ΔND_, the 27 amino acid (aa122-148) deletion of ND domain and Rev_L95D_, described in the materials and methods. WB analysis of Mat (B) or Gag (D) expression levels when a Mat expression plasmid or a Rev-dependent EIAV Gag/Pol expression construct (pGP-RRE) was transfected into HEK293T cells with either an empty vector VR1012 (as a negative control) or a series of Rev expression plasmids, as indicated. Distribution of cytoplasmic and nuclear mat (C) or gag (E) mRNAs in HEK293T cells at 36 pht was analyzed using qPCR when Mat or pGP-RRE expression plasmids were transfected into HEK293T cells with a series of Rev expression plasmids. All of the experiments were performed three times, and a representative result is shown.

In addition, a Rev-dependent EIAV Gag-Pol expression vector, pGP-RRE (described in the materials and methods), was transfected into HEK293T cells in the presence of Rev_WT_ or different Rev mutants. The levels of Gag protein were assessed using WB and the distribution of gag mRNA in the cytoplasm and nucleus was analyzed using qPCR. No Gag protein was detected in the absence of Rev or in the presence of Rev_L95D_ (Fig 4D, lanes 1 and 7). Compared to those in the presence of Rev_WT_, very little Gag protein was detected in the presence of Rev_mNES_, Rev_mRBD_ or Rev_mNLS_ (Fig 4D, lanes 3, 4 and 5), but none of Gag proteins expression level showed remarkable changes in the presence of Rev_ΔND_ compared with in the presence of Rev_WT_ (Fig 4D, lane 6). The qPCR analysis also showed that gag mRNAs were mainly distributed in the cytoplasm in the presence of Rev_WT_ or Rev_ΔND_, while the presence of other Rev mutants gave results similar to those obtained in absence of Rev, with the gag mRNAs being mainly distributed in the nucleus (Fig 4E). Consistent with the results of previous studies [30], these results also indicate that in addition to ND, other functional domains of Rev play an important role in mediating the nuclear export of gag-pol mRNAs.

Taken together, these results show that the expression levels of Mat or Gag protein are related to the accumulation of the mRNAs in the cytoplasm. Interestingly, we found that the export of mat and gag mRNAs from the nucleus relied on different function domains of Rev, indicating that Rev mediates the expression of Gag and Mat through different mechanisms.

### The first exon of Mat cannot replace the known RRE to mediate EIAV Gag/Pol expression

As described above, the first exon of mat can be used as an RNA export element to bind to Rev to mediate the expression of Mat. Therefore, in order to test whether the mat-exon1 can replace the known RRE to mediate the expression of Gag/Pol, a series of EIAV Gag/Pol constructs was prepared: the pGP contained only the EIAV gag/plo-coding sequences; pGP-RRE, pGP-MA and pGP-3×MA were based on pGP with an EIAV RRE (22), or one copy of mat-exon1, or three copies of mat-exon1 inserted respectively downstream of the gag/pol coding sequence (Fig 5A). HEK293T cells were transfected with these constructs in the presence or absence of Rev, and the levels of Gag protein in the cells were analyzed using WB. Gag expression could only be detected in cells transfected with pGP-RRE in the presence of Rev (Fig 5B, lane 4), while we could not detect Gag expression in cells transfected with other constructs with or without Rev. Therefore, these results indicate that the Mat first exon sequence within the Gag/pol-coding sequence was unable to mediate the expression of Gag/Pol under the effect of Rev (Fig 5B, lane 2), and the first exon of mat cannot replace RRE to mediate the expression of Gag/Pol (Fig 5B, lanes 6 and 8). Taken together, these results show that single or multimer copies of mat-exon1 cannot replace RRE function in the EIAV Gag/Pol expression vector.

**Fig 5.**
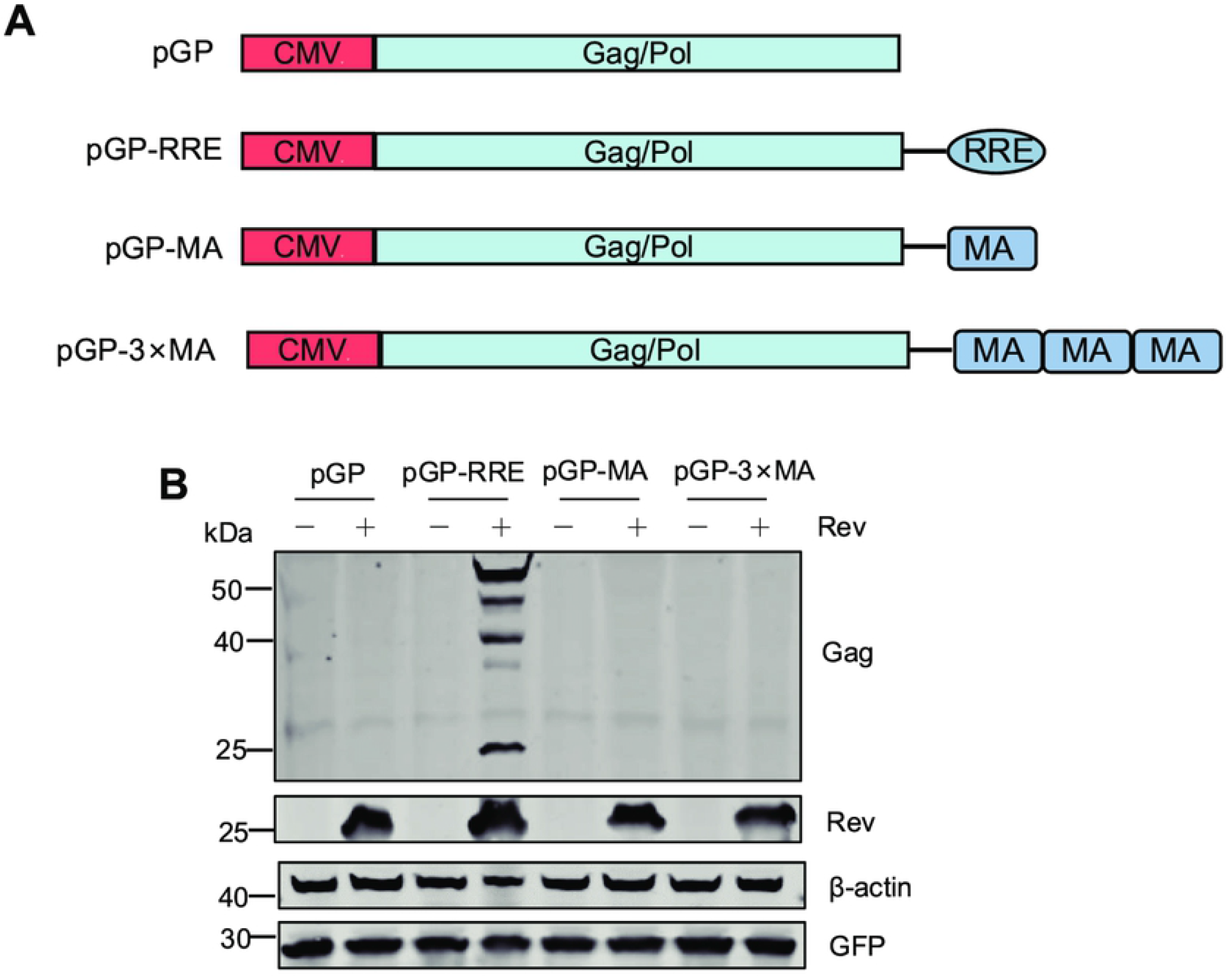
Functional identification of Mat first exon used as an RNA export element. (A) Schematic diagram of a series of plasmids used for the identification of the function of the RNA export element. pGP: contains only EIAV gag/plo-coding sequences; pGP-RRE: with EIAV RRE inserted downstream of the gag/pol coding sequence; pGP-MA: with one copy of mat-exon1(MA) inserted downstream of the gag/pol-coding sequence; pGP-3×MA with three copies of mat-exon1(MA) inserted downstream of the gag/pol-coding sequence. (B) WB analysis of indicated plasmids cotransfected with Rev or empty vector. All of the experiments were performed three times, and a representative result is shown.

## Discussion

Lentivirus transcription takes place in the nucleus, and the various transcripts produced by transcription usually undergo nuclear export via two pathways for gene expression [8]. In the case of HIV-1, nuclear export of the completely spliced transcripts is mediated by cellular mRNA transporter proteins, while nuclear export of the unspliced full-length and incompletely spliced transcripts is mediated by Rev through binding to RRE via the CRM1 pathway. In EIAV, only one RRE has been found to date, and this is located in the Env-coding region [22, 32]. In this report, we found a novel fully-spliced transcript in EIAV, encoding a 20kDa protein, and named it Mat. Interestingly, we demonstrated that the expression of Mat depends on Rev, that Rev can increase the nuclear export of Mat mRNAs, and that Rev can specifically bind to Mat mRNAs. Furthermore, the first exon of Mat mRNAs (Mat-exon1 mRNA) has been identified as a key region for Rev-mediated nuclear export. These results indicate that, in addition to the reported previously RRE located in the Env-coding region in EIAV, there is a second RRE in the Mat coding region. These findings have updated our knowledge of EIAV genome structure and its gene transcription pattern, as illustrated in Fig 6.

**Fig 6.**
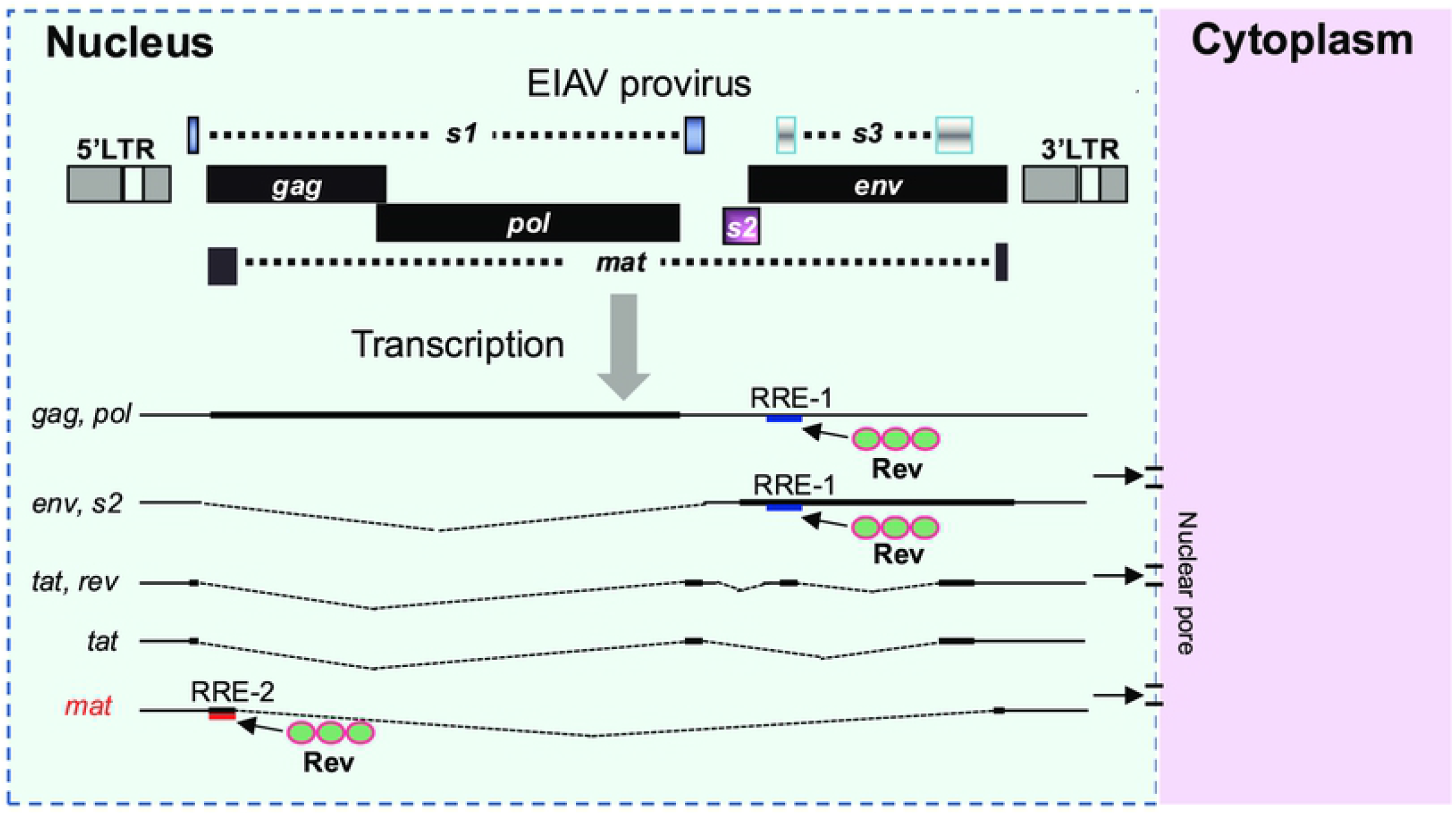
The EIAV proviral genome structure and transcriptional patterns. The previously identified Rev-response element (RRE) was named RRE-1 (blue line), and is located in the env-coding region. The unspliced full-length mRNA transcripts encoding Gag and Pol proteins and an incompletely spliced transcript encoding Env and S2 proteins are exported to the cytoplasm in a pathway mediated by the binding of Rev to RRE-1. The transcripts encoding Tat or Rev protein are exported to the cytoplasm via an endogenous cellular pathway used by cellular mRNAs independent of Rev. The transcripts encoding Mat protein are exported to the cytoplasm via a pathway mediated by the binding of Rev to the first exon of Mat mRNA. The first exon of mat therefore has the function of RRE, and is named here RRE-2 (red line).

In lentiviruses, alternative splicing is a common phenomenon that promotes the production of various transcripts encoding different viral proteins. Previous studies have shown that five major transcripts are produced from the EIAV genome [15, 16], encoding three structural proteins, three accessory proteins, and a protein with unknown function, Ttm. Except for the unspliced full-length transcripts, all spliced transcripts share a common 5’-splice donor site. We recently found an unreported transcript in EIAV encoding a protein that antagonizes the antiviral activity of the host protein Tetherin (manuscripts, under review). In this study, we once again report a novel EIAV transcript, which is produced by a single splicing event, with a 5’-splice donor site that is different from other reported splicing transcripts. This suggests that EIAV alternative splicing patterns are complex. Importantly, we have confirmed that this transcript encodes a novel viral protein, here designated Mat. Although the biological function of Mat is unclear at present, our research results have further updated our knowledge of the genome structure of EIAV and of its patterns of gene transcription.

Cis-acting repressing sequence (CRS) is a type of RNA sequence element that regulates gene expression by inhibiting the nucelar export of mRNAs [33]. It is widely present in a variety of viruses including HIV-1 [34–36], EIAV [37], HBV [38] and HPV [39], as well as in cellular transcripts. Although no uniformly recognizable sequence commonality was detected in CRS, it has been determined that a variety of host proteins can specifically bind to CRS and restrict RNA in the nucleus [40–42]. In HIV-1, several CRS have been identified in multiple regions such as gag, pol and env genes and it has been demonstrated that these inhibitory sequences can prevent nuclear export of unspliced and partially spliced transcripts, but the supply of Rev overcomes this inhibition [34, 36, 43]. A previous study by Rosin-Arbesfeld et al. also found that a CRS exists in the center of the EIAV genome (the C-terminus of the Pol coding region), and also demonstrated that this CRS can inhibit the nuclear export of viral transcripts [37]. In this study, we found that the mat or mat-exon1gene constructs showed poor expression and cytoplasmic RNA accumulation in the absence of Rev, suggesting the existence of a potential CRS in the first exon of the mat gene. Interestingly, a CRS has been identified at the N-terminus of the HIV-1 gag gene in a previous study [35]. In fact, the first exon of the mat gene is also located at the N-terminus of the EIAV gag gene. However, the gag genes of HIV-1 and EIAV lack sequence homology. Importantly, the expression of Mat or Mat-exon1 was rescued by co-expression with Rev, indicating that Rev may overcome the inhibitory effect of the mat-exon1 gene, which also suggested that mat-exon1 sequences contain an element enabling Rev-mediated nuclear export, such as an RRE. These results suggest that both a CRS and RRE may exist simultaneously in the mat-exon1, or the two may overlap with each other. Notably, the overlap of a CRS and RRE has been documented HIV-1 [43].

Interestingly, we found that the expression of Mat is dependent on Rev. Rev is an accessory protein necessary for lentiviral replication, and its biological function is to mediate the transport of the unspliced and incompletely spliced viral mRNAs from the nucleus to the cytoplasm. Mechanistically, Rev binds to an RRE located within the Env-coding region, then polymerization occurs, and the polymerized protein facilitates the export of the Rev-RNA complex into the cytoplasm through a CRM1-dependent pathway. We found that Rev could specifically bind to mat mRNAs, and that mutation of the Rev multimerization site disrupted its ability to mediate Mat expression. In addition, the use of the CRM1 inhibitor LMB significantly decreased Mat expression levels. These results demonstrate that the expression of Mat depends on the nuclear export activity of Rev. Previous studies have identified the RRE of EIAV that is located in a 555 nt region near the 5’ end of env gene [22, 32]. In this study, we found that Rev could bind to Mat-exon1 mRNAs and mediate their nuclear export, and this phenomenon has been verified in 5 other EIAV strains. This indicates that the first exon (393 nt) of mat mRNAs is the second known RRE of EIAV. We named it RRE-2, with the reported RRE located in the Env coding region defined as RRE-1. In HIV-1, an additional RRE was also found in the packaging signal located in 5’-untranslated region (5’UTR) of the genome [44, 45]. Here this HIV-1 RRE is termed 5’RRE. Mutation of 5’RRE led to reduced nuclear export of viral genomic RNA (gRNA) and decreased gRNA packaging efficiency but did not affect the production of viral proteins and the export of viral particles, however, replication of the virus was severely disrupted [45]. Rem is an accessory protein encoded by mouse mammary tumor virus (MMTV) and is a homolog of the Rev protein in lentivirus. MMTV, a member of the genus Betaretrovirus, harbors two Rem-responsive elements (RmREs): a 5’RmRE located at the 5’UTR of gRNA that is required for nuclear export of unspliced RNA, and a 3’ RmRE located at env-U3 junction region that is needed for expression of both unspliced and spliced transcripts [46]. Therefore, EIAV is different from HIV-1 and MMTV in that its two RREs are located in the coding region sequence. We noticed that RRE-2 plays the nuclear export function of Rev-RRE only in the context of the mat transcript, but not in the context of gag/pol mRNA. However, RRE-1 is able to mediate transport of both Env-encoding transcripts and Gag/Pol-encoding transcripts. Possible explanations for this difference include that the nuclear export ability of Rev-RRE-2 complex is weaker than that of Rev-RRE-1, or that the structure of RRE-2 may be more susceptible to interference from adjacent sequences.

We also observed that the nuclear export functions of Rev-RRE-1 and Rev-RRE-2 are dependent on different amino acid sites and regions of Rev. This suggests that the molecular mechanisms by which Rev mediates nuclear export of diverse target RNAs may differ, although to date no similar phenomenon has been reported in lentiviruses. We speculate that this may be related to the three-dimensional structure of Rev or to the involvement of host factors [20, 27, 47].

In summary, we identified a novel protein, Mat, from EIAV, and demonstrated that the the expression of Mat depends on Rev, and that the first exon of mat mRNA acts as a Rev binding element to bind Rev and mediate Mat expression via CRM1 pathway. These findings update the EIAV genome structure and highlight the diversification of post-transcriptional regulation patterns in EIAV, and provides important insights into the Rev-dependent nuclear export of completely spliced transcripts in lentiviruses.

## Materials and Methods

### Plasmids

The Mat expression plasmid, pcDNA-Mat-HA, was constructed by inserting the mat gene sequence (GenBank accession numbers: OM417133) into a pcDNA3.1 (+) vector and fusing HA-tags to the C-terminal. To constructed a codon-optimized Mat expression plasmid, pcDNA-optMat-HA, we cloned the sequence corresponding to mat-exon1 from the codon-optimized Gag plasmid and inserted it before the mat-exon2 sequence. The plasmid pcDNA-Mat-exon1_FDDV_-HA was constructed by cloning the first exon of mat gene from pCMV3-8 [48] and inserting it into pcDNA3.1-HA. The other 5 Mat-exon1 expression plasmids were synthesized according to the following EIAV strains genome sequence: EIAV_UK_ (GenBank accession numbers: AF016316.1; nt450-836), EIAV_Miyazaki-2011-A_ (GenBank accession numbers: JX003263.1; nt450-836), EIAV_Cornwall_ (GenBank accession numbers: MH580898.1; nt231-617), EIAV_DE_ (GenBank accession numbers: KM247554.1; nt114-500) and EIAV_NC-Dagmar_ (GenBank accession numbers: MH820164.1; nt100-486). The Rev expression plasmid, VR-Rev-Flag, was constructed by inserting the EIAV Rev gene sequence into a VR1012 vector and fusing Flag-tags to the C-terminal. VR-Rev_HIV_-Flag was constructed by substituting the HIV-1 Rev gene sequence (Genbank accession numbers: AF324493.2) into VR-Rev-Flag. A series of Rev-mutant plasmids were generated using site-directed mutagenesis on the template plasmid VR-Rev-Flag. The complete EIAV gag-pol gene was PCR amplified from the molecular clone pCMV3-8 and inserted into the pEGFP-N1 vector (Clontech, USA) to construct the plasmid pGP. The Rev-responsive element (RRE) was cloned from pCMV3-8, and inserted into the EcoR1 and Not1 sites of pGP to construct the pGP-RRE plasmid. The mat-exon1 sequence was cloned as a EcoR1-Not1 fragment into pGP, which obtained plasmids containing a single mat-exon1(pGP-MA) as well as multimer copies of the mat-exon1(pGP-3MA). All constructed plasmids were verified by sequencing.

### Cell culture and transfection

Human embryonic kidney epithelium 293T (HEK293T) cells and Hela cells were cultured in DMEM-high glucose (Sigma-Aldrich, USA) supplemented with 10% fetal bovine serum (Sigma-Aldrich, USA) and 1% penicillin-streptomycin (Gibco, USA). Equine monocyte-derived macrophages (eMDM) cells were isolated from fresh heparinized horse whole blood as previously described [49], and cultured in RPMI 1640 (Sigma-Aldrich, USA) supplemented with 20% donor equine serum (HyClone, USA) and 40% newborn bovine serum (Ausbian, Australia). Fetal donkey dermal (FDD) cell cultures were maintained in Minimal Essential Medium (α-MEM, Gibco, USA) supplemented with 10% fetal bovine serum (Sigma-Aldrich, USA) and 1% penicillin-streptomycin (Gibco, USA) [50]. All cells were maintained in a humidified incubator at 37°C containing 5% CO_2_.

Cells were transfected with the indicated plasmids using PolyJet DNA transfection reagent (SignaGen, USA), following the manufacturer’s instructions. EGFP-expression construct (pEGFP-N1) was added to all transfections to monitor transfection efficiency. All transfections were repeated at least three times.

### Amplification and identification of mat

For the amplification of mat, polyadenylated RNAs were prepared from frozen horse tissues infected with EIAV_LN40_ using a Trizol method [51]. The tissues were homogenized in liquid nitrogen, and 1ml of Trizol regent was used to incubate each sample. RNAs were precipitated with 0.5ml isopropyl alcohol and then washing with 1ml 75% ethanol to obtain purified total RNAs. For further identification of the mat transcript, an EIAV virulent strain (EIAV_DLV36_) or vaccine strain (EIAV_DLV121_) at equivalent titer was used to infect eMDM cells cultured in T25 flasks [49]. At 4 days post-infection, cells were harvested and total RNAs were extracted using a Bio-fast simply P RNA extraction kit (Bioer, China).

About 1μg of RNAs were reverse transcribed into cDNAs using a PrimeScript RT Reagent Kit with gDNA Eraser (Takara, Japan) following the manufacturer’s protocol. EIAV transcripts were amplified with Kod FX Neo polymerase (Toyobo, Japan) in polymerase chain reactions (PCR). Nested primer pairs included: p399 (5’-GGACAGCAGAGGAGAACTTACAG-3’; forward primer), p8144 (5’-AAGGGACTCAGACCGCAGAATCT-3’; reverse primer) / p8081 (5’-TAAAAACAGGAASTTAACGCGTCAC-3’; reverse primer) were used to amplify EIAV transcripts. Nested primer pairs: p555 (5’-TCAAAAGCTAACTAATGGTAAC-3’; forward primer) / p7878 (5’-ATTGAGGCATTGTTACATGAGA-3’; forward primer), p643 (5’-CAATTAAGGGACGTCATTCCAT-3’; reverse primer) / p7855 (5’-GTAGCTGGATTTAGCATTTTCC-3’; reverse primer) were used to identify mat-specific transcripts. The amplification conditions were as follows: 2min at 98°C; 35 cycles of 30s at 98°C, 30s at 56°C and 10-30s at 72°C; and a final extension step of 10min at 72°C. PCR products were separated using electrophoresis on 1% agarose gels in 1×Tris -acetate-EDTA, and were visualized by staining with ethidium bromide. The amplified fragments were purified using a gel extraction kit (Vazyme, China) and cloned into a pMD-18T vector (Takara, Japan).

### Western blotting

The expression of the indicated plasmids was analyzed using western blotting as previously described [52]. In brief, cells were harvested 48 hpt and resuspended in lysis buffer (150mM Tris-HCl [PH 7.6], 50mM NaCl, 5mM EDTA, and 1%Triton X-100). Total protein extracts were separated by SDS-PAGE (Genscript, USA) and then blotted to NC membranes (Millipore, Germany). The membrane was subsequently blocked for 2 h with 5% skim milk (Biosharp, China) in PBS at room temperature. PBS containing 5% skim milk was used to dilute primary and secondary antibodies. After washing, the membrane was analyzed on the LI-COR Odyssey Imaging System (LI-COR, USA). All experiments were performed at least in triplicate.

### Antibodies

Mat proteins in eMDM cells infected with EIAV_DLV121_ and EIAV_DLV36_ were detected using a mouse anti-Mat monoclonal antibody (mAb), which was produced by immunization of mice with a Keyhole Limpet Hemocyanin (KLH) carrier protein containing a polypeptide corresponding to the C-terminal 15 amino-acids (ENAKSSYISCNNASI) of Mat. Gag proteins were analyzed using a mouse anti-p26 mAb (9H8), kept in our lab [53]. The primary antibodies purchased from commercial sources are listed below: mouse anti-HA mAb (Sigma, USA), mouse anti-FLAG mAb (Sigma, USA), mouse anti-GFP mouse mAb (Affinity Bioscience, USA), mouse anti-LaminB1 mAb (Proteintech, China), mouse anti-α tubulin (Abcam, UK), and mouse anti-β actin mAb (Sigma, USA). Goat anti-mouse IRD800-conjugated mAb (Sigma, USA) was used as secondary antibody for western blotting.

### Cell fractionation and quantitative PCR analysis

For the preparation of nuclear and cytoplasmic RNAs, HEK293T cells were harvested 36 hpt and purified using a PARIS protein and RNA isolation kit (Invitrogen, Thermo Fisher Scientific, USA) according to the manufacturer’s protocols. Equal volume RNAs were subjected to reverse transcription using the PrimeScript RT Reagent Kit with gDNA Eraser (Takara, Japan). Distribution of mat mRNAs or mat-exon1 mRNAs in transfected cells was assessed using SBYR-Green (Takara, Japan)-based quantitative PCR analysis on Aligent Mx3005P. The primers used are as follows: Mat-forward (5’-AAGGGACGTCATTCCATTGTT-3’) and Mat-reverse (5’-ATTTGTAAGCCCATCTTAACG-3’); GFP-forward (5’-GCAAGCTGACCCTGAAGTTCATC-3’) and GFP-forward (5’-GTCTTGTAGTTGCCGTCGTCCTT-3’) Mat-exon1-forward (5’-AGTTAGAGAAGGTGACGGTAC-3’) and Mat-exon1-reverse (5’-CAACAATGGAATGACGTCCCT-3’); Gag-forward (5’-CGATGCCAAATCCTCCATTAG-3’) and Gag-reverse (5’-CTGATCAAAAGCAGGTTCCATCT-3’).

### *In vitro* transcription and RNA-pull down assay

Template plasmids were digested using Not1 (Thermo Scientific, USA) to generate linearized DNAs, which were subsequently transcribed to produce copies of the mRNAs using an *in vitro* transcription kit (New England Biolabs, UK) according to the manufacturer’s protocol. The equimolar mRNAs were labeled using biotin with an RNA 3’ End Desthiobiotinylation Kit (Thermo scientific, USA). The labeled mRNAs were then captured using streptavidin magnetic beads (Thermo scientific, USA). After washing three times with Protein-RNA Binding Buffer (20mM Tris [pH 7.5], 50mM NaCl, 2mM MgCl_2_, 0.1% Tween-20 Detergent), 100μg of the total protein extracted from transfected cells was added to the RNA-bound beads and incubated for 1h on a roller at 4°C. Beads were then washed five times with Washing Buffer (20mM Tris [pH 7.5], 10mM NaCl, 0.1% Tween-20 Detergent), and the beads containing RNA-binding protein complexes were harvested and analyzed with Western blotting.

## Acknowledgements

The authors thank Yandong Tang for their constructive suggestions in this study. This study was supported by grants from the National Natural Science Foundation of China (32170169 to X. W, 32172831 and 31672578 to XF.W), the Natural Science Foundation of Heilongjiang Province of China (C2017076 to XF.W), and postdoctoral scientific research developmental fund of Heilongjiang Province (LBH-Q20059 to XF.W).

## Notes

### Competing Interest Statement

The authors have declared no competing interest.

